# Olfactory host entry supports herpesvirus recombination

**DOI:** 10.1101/2020.08.24.264903

**Authors:** Wanxiaojie Xie, Kimberley Bruce, Helen E. Farrell, Philip G. Stevenson

## Abstract

Herpesvirus genomes record abundant recombination. Its impact on infection remains ill-defined. When co-infecting mice by the natural olfactory route, individually incapacitated Murid Herpesvirus-4 (MuHV-4) mutants routinely recombined to restore normal host colonization. Lung infection rescued much less well. Murine cytomegalovirus mutants deficient in salivary gland colonization also showed rescue via the nose but not the lungs. As nose and lung infections show similar spread, efficient recombination seemed specific to olfactory entry. Rescue of replication-deficient MuHV-4 implied co-infection of the first encountered cells, and this worked also with asynchronous inoculation, suggesting that latent virus could lie in wait for later reactivation. Inhaled MuHV-4 is commonly caught on respiratory mucus, which epithelial cilia push back towards the olfactory surface, and infection was correspondingly frequent at the anterior olfactory edge. Thus olfactory entry provides a general means for herpesviruses to meet.

**Author summary:** Inter-strain recombination allows viruses to optimise infection in diverse hosts. Many herpesviruses show past recombination. Yet they are ancient pathogens, so this past may be remote and recombination rare. Diverse herpesviruses enter new hosts via olfactory cells. We show that such entry routinely allows recombination between co-infecting virus strains, even when one strain cannot spread. Recombination was contrastingly rare after lung infection. Thus, entry via olfactory cells specifically supports frequent herpesvirus recombination.

## Introduction

Outbred hosts subject viruses to serial changes in selection. Herpesviruses, with proof-reading polymerases, are slow to make new mutations [1]. Nonetheless some human cytomegalovirus (HCMV) and Herpes simplex virus type 1 (HSV-1) genes display marked diversity, including disruption [2, 3]. The latency genes of Epstein-Barr virus (EBV) also vary greatly [4]. Ancient diversity plus present stability gives herpesvirus genes allelic forms, which interact with host alleles to influence infection outcomes. For example the murine cytomegalovirus m157 evolved to inhibit NK cells [5], but in Ly49H^+^ mice delivers instead disadvantageous activation [6, 7].

Herpesvirus strains often co-infect [2–4], and genome sequences show that co-infections must have met to make recombinants [8–11]. However each infection is dilute, and immunogenicity creates inter-strain competition. Longitudinal analysis has identified likely instances of HCMV recombination [12], but in immunodeficient hosts with high viral loads and susceptibility to re-infection. While EBV co-infection is common in infectious mononucleosis [13], that is 1-3 months after entering an immunocompetent host [14], serial strain typing has been limited to single loci. Thus it is unclear whether divergent strains co-infect cells frequently enough to form functional gene pools, or whether recombination rarely alters outcomes.

Recombination seems most likely to result in early infection, when viral loads are largest. Early human herpesvirus infections are hard to sample, so their routes and recombination opportunities remain obscure. Experimental infections are more accessible. Yet most employ invasive inoculations, via virus injection or aspiration under anaesthesia [15, 16], which may bypass (or make) bottle-necks for recombination. The prominent presence of EBV in tonsils during infectious mononucleosis led to assuming that γ-herpesviruses enter orally [17, 18]. However EBV DNA appears in blood before saliva [19], arguing that tonsillar infection is host exit, not entry. This result was anticipated by analysis of the related Murid Herpesvirus-4 (MuHV-4), which found orally fed virions were non-infectious unless they reached the respiratory tract [20], and submucosal B cell colonisation was subsequent to systemic spread [21]. MuHV-4 enters minimally manipulated mice via the olfactory epithelium [22]. The only natural non-olfactory uptake known is genital [23]. Olfactory infection spreads to lymph nodes, via dendritic cells [24], then to the spleen and beyond via B cells [25].

While olfactory entry remains unproven for EBV, it is evident for MCMV, including spontaneous transmission [26], and for HSV-1 [27]. HCMV has an olfactory receptor [28]. MuHV-4, MCMV and HSV-1 diverged hundreds of millions of years ago [29], so their shared olfactory entry suggests that many mammalian herpesviruses use the same route. We tested with MuHV-4 whether it offers an opportunity for co-infecting viruses to recombine.

## Results

### Nasal infection with latency-deficient MuHV-4 mutants

γ-herpesviruses colonize their hosts lytically and by latency-associated lymphoproliferation. MuHV-4 mutants lacking stable latency have been made by over-expressing the ORF50 viral transactivator [30, 31], or by disrupting ORF73 episome maintenance [32, 33]. M50 MuHV-4 has an MCMV IE1 promoter fragment inserted in the 5’ untranslated region of ORF50, deregulating its transcription [30]. ORF73FS MuHV-4 has a frameshift mutation in ORF73 [32] (Fig.1a). Both mutants replicate lytically in the lungs after a 30μl inoculation under anaesthesia, but poorly colonize lymphoid tissue. Nasal inoculation (5μl without anaesthesia) similarly led to local lytic spread but signficantly less lymph node infection than wildtype, with none detectable in spleens at day 18 (Fig.1b).

**Figure 1.**
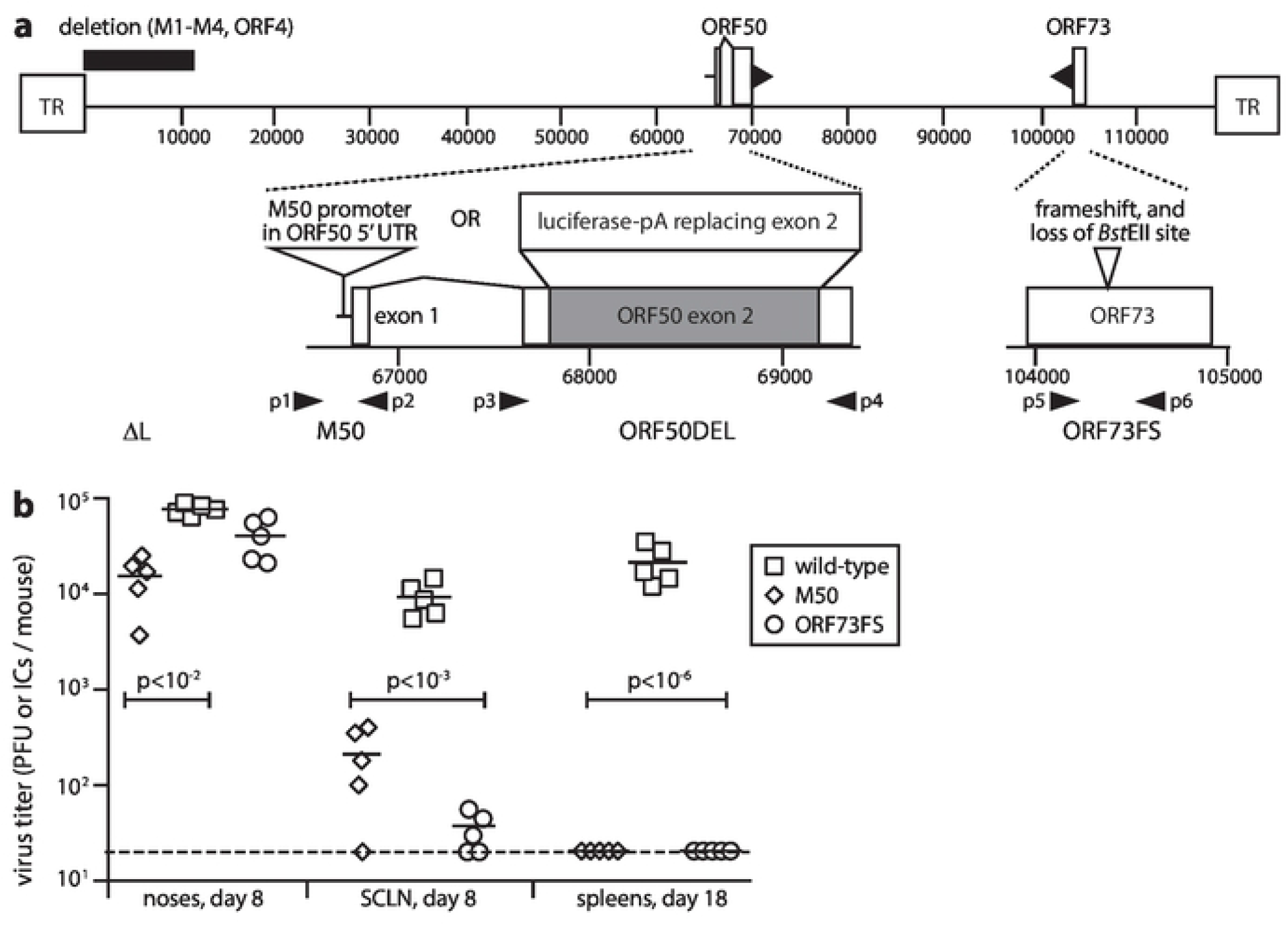
Nasal infection by MuHV-4 mutants lacking normal latency. **a.** A schematic sketch of MuHV-4 mutations shows the linear genome flanked by terminal repeats (TR), with expanded views below. Replication-deficient ORF50DEL MuHV-4 has most of ORF50 exon 2 replaced by luciferase plus a polyadenylation site. ORF73FS MuHV-4 has a frameshift in ORF73, which encodes the viral episome maintenance protein. The ΔL virus mutation additionally deletes 11kb from the genome left end, removing ORFs M1-M4. M50 MuHV-4 has an MCMV IE1 promoter inserted in the 5’ untranslated region of ORF50. p1-p6 show the locations of primers used to identify the ORF50DEL, M50 and ORF73FS mutations. **b.** C57BL/6 mice were infected nasally (10^5^ p.f.u. in 5μl without anaesthesia) with wild-type, M50 or ORF73FS virus. Noses were titered by plaque assay. Superficial cervical lymph nodes (SCLN) and spleens were titered by infectious centre assay. Symbols show individual mice, bars show means. The dashed line shows the detection limit. Significant differences in virus recovery relative to wild-type are shown.

### Recombinational rescue of latency-deficient mutants after nasal co-infection

To test rescue by recombination, we co-infected mice nasally with ORF73FS and M50 MuHV-4, then 25 days later determined latency by infectious centre assay (Fig.2). 11/12 co-infected mice showed significant splenomegaly (Fig.2a) and splenic infection (Fig.2b). No singly infected mice did so - the M50 [30] and ORF73FS mutations [32, 34] have consistently not shown spontaneous reversion. Co-infection alone might complement ORF73 deficiency, but ORF50 over-production should have a dominant detrimental impact on latency. Therefore recombinational rescue seemed more likely. This was confirmed by PCR analysis of viruses cloned from splenic infectious centres. Clones from 4/4 co-infected mice showed M50 and ORF73 PCR products of wild-type size (Fig.2c, 2d), and their DNA sequences exactly matched wild-type.

**Figure 2.**
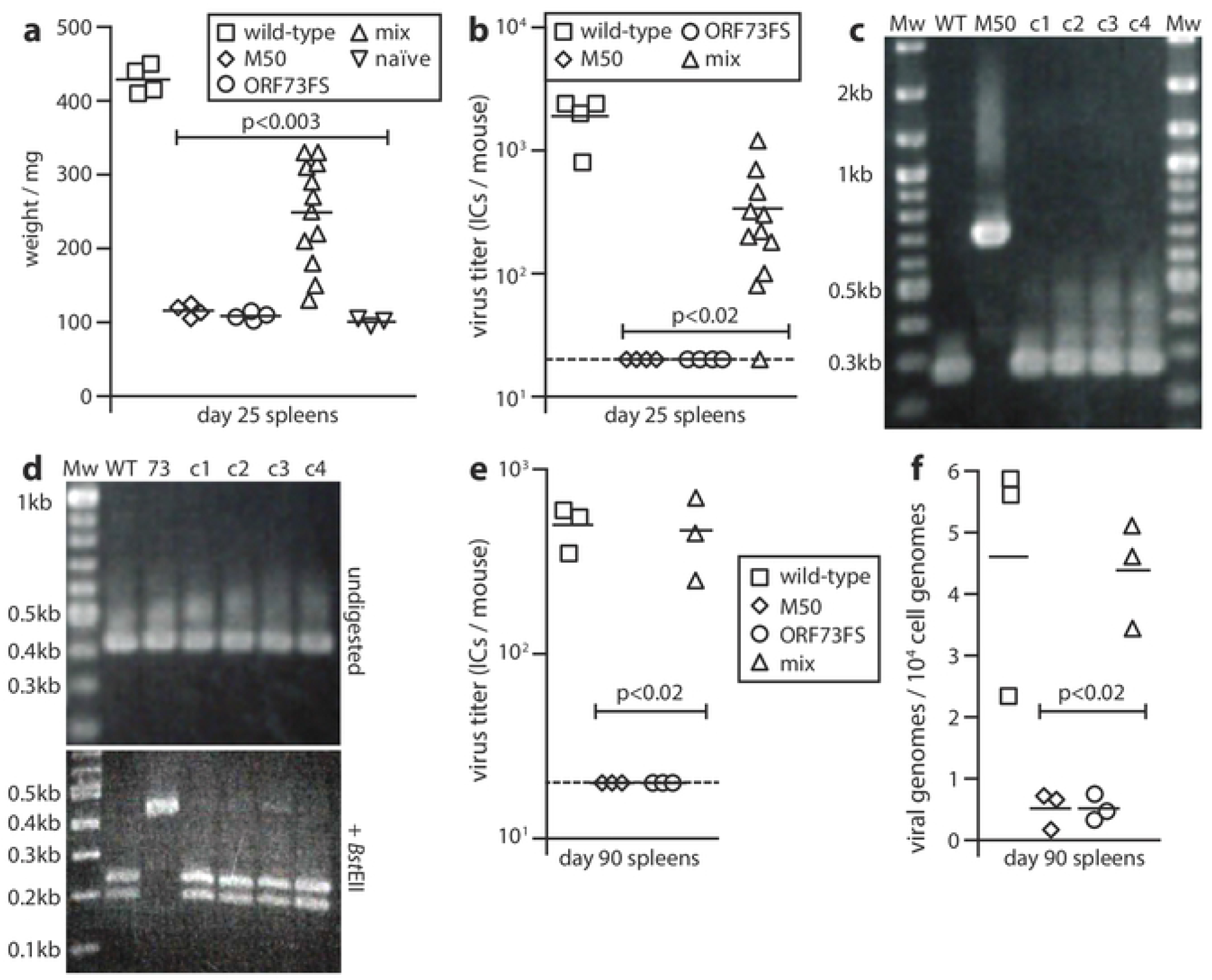
Recovery of latency after nasal co-infection with latency-deficient MuHV-4 mutants. **a.** C57BL/6 mice were infected nasally (10^5^ p.f.u.) with wild-type, M50 or ORF73FS virus, or a 1:1 M50 / ORF73FS mix. Weigts of spleens 25 days later are shown, compared to naive mice, with significant splenomegaly for the wild-type and mixed infections. Symbols show individual mice, bars show means. **b.** The spleens in **a** were titered for latent virus by infectious centre assay. Symbols show individuals, bars show means. The dashed line shows the lower limit of assay detection. Mixed M50 / ORF73FS infection yielded significantly more virus than either single infection. No preformed infectious virus was recovered by parallel titer of freeze-thawed spleen samples. **c.** Viruses cloned (c1-c4) from spleens of mixed infection mice were genotyped by PCR across the M50 insertion site (primers p1 and p2 in Fig.1a). PCR products were resolved by agarose gel electrophoresis and stained with ethidium-bromide. WT and M50 are wild-type and M50 input viruses. Mw = molecular weight markers. Sequencing of the main (268bp) product of cloned virus DNA confirmed identity with the wild-type. **d.** DNA from the cloned viruses in **c** was genotyped by PCR across the ORF73 frameshift (primers p5 and p6 in Fig.1a). PCR products were digested or not with *Bst*EII, resolved by agarose gel electrophoresis, and stained with ethidium-bromide. WT and 73 are wild-type and ORF73FS input viruses. As the viruses were cloned prior to analysis, we interpret the minor residual 0.43kb product for *Bst*EII-digested c3 DNA as incomplete digestion. DNA sequencing of the main (428bp) undigested product of the cloned viruses confirmed identity with the wild-type. **e.** C57BL/6 mice were infected as in **a**. 3 months later, latent virus was detected by infectious centre assay. M50 / ORF73FS co-infection yielded significantly more virus than either single infection, and was indistinguishable from wild-type. Symbols show individual mice, bars show means. The dashed line shows the detection limit. **f.** The spleens in **e** were assayed for viral genomes by quantitative PCR of extracted DNA. Viral copies are expressed relative to cellular β-actin copies amplified in parallel. M50 / ORF73FS co-infection yielded significantly more viral genomes than either single infection, and was indistinguishable from wild-type.

That recombinants showed less early splenomegaly and latency than wild-type was unsurprising: even cells co-infected with MuHV-4 *in vitro* yield only a few percent recombinants (PGS, unpublished data), so they would inevitably initiate at low dose. After 3 months, the splenic loads of co-infected mice were indistinguishable from wild-type, by infectious centre assay (Fig.2e) and by quantitative PCR of viral DNA (Fig.2f). Thus, while the M50 and ORF73FS viruses vaccinate against a later wild-type challenge [35, 36], immunity did not impair the outgrowth of recombinant infection.

### Dual recombinational rescue

Linear genomes can repair single mutations by one recombination. Herpesvirus genome ends vary more than their middles, and one recombination would allow co-infecting viruses to exchange left or right end ends. However in more detail, the genome core comprises conserved blocks, between which lie more varied loci. To exchange these while retaining left and right ends would require two recombinations. To simulate this setting, we co-infected mice with M50 MuHV-4 and an ORF73FS mutant that also lacked 10kb from its left end (ORF73FSΔL) [37] (Fig.1a). The ΔL mutation deletes ORFs M1-M4, and independently impairs the establishment of a normal latent load [38]. Nasal co-infection nonetheless gave rescue (Fig.3), implying recombination both between ORFs 50 and 73, and between ORF50 and ΔL (see Fig.1a). Thus there was no barrier to more complex categories of recombinational rescue.

**Figure 3.**
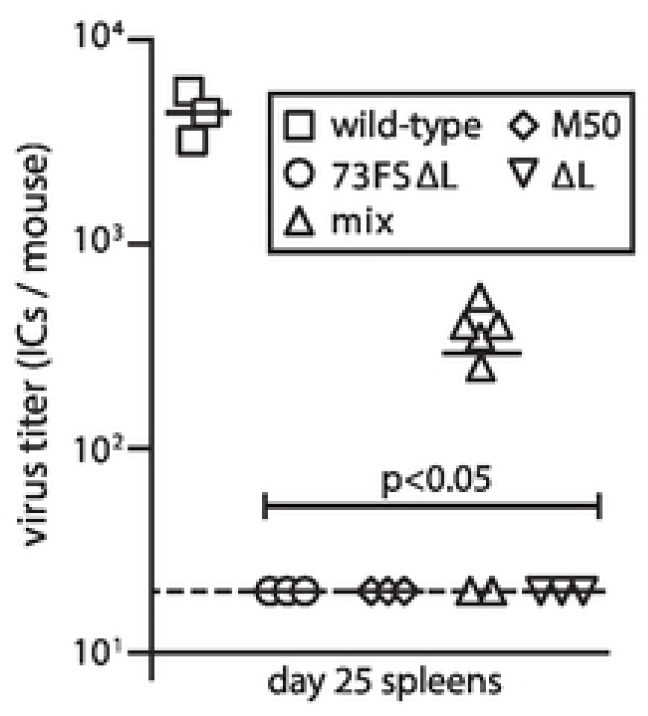
Recovery of latency after co-infection by M50 and ORF73FSΔL mutants. C57BL/6 mice were infected nasally (10^5^ p.f.u.) with wild-type, M50 or ORF73FSΔL virus, or a 1:1 M50 / ORF73FSΔL mix. We also infected mice with a control ΔL single mutant. Spleens were infectious centre-assayed for latent infection 25 days later. Symbols show individual mice, bars show means. The dashed line shows the detection limit. M50 / ORF73FSΔL co-infection yielded significantly more latency than either virus alone. No preformed infectious virus was recovered by parallel titer of freeze-thawed spleen samples.

### Lung co-infection with latency-deficient MuHV-4 mutants

To determine whether MuHV-4 co-infection generally provides genetic rescue, or whether this a particular property of olfactory entry, we tested lung inoculation, a widely used immunological model. We gave mice M50 and ORF73FS MuHV-4 as before but in 30μl under anaesthesia. As alert mice retain little free fluid in their upper airways [39], lung inoculation is more efficient than nasal. Therefore we gave 10^4^ rather than 10^5^ p.f.u. of virus. We assayed spleens for latent load 17 days later (Fig.4a) - as lung infection is more extensive than olfactory, it drives faster down-stream spread [20]. Low levels of latent infection were seen in the spleens of 3/6 mice.

**Figure 4.**
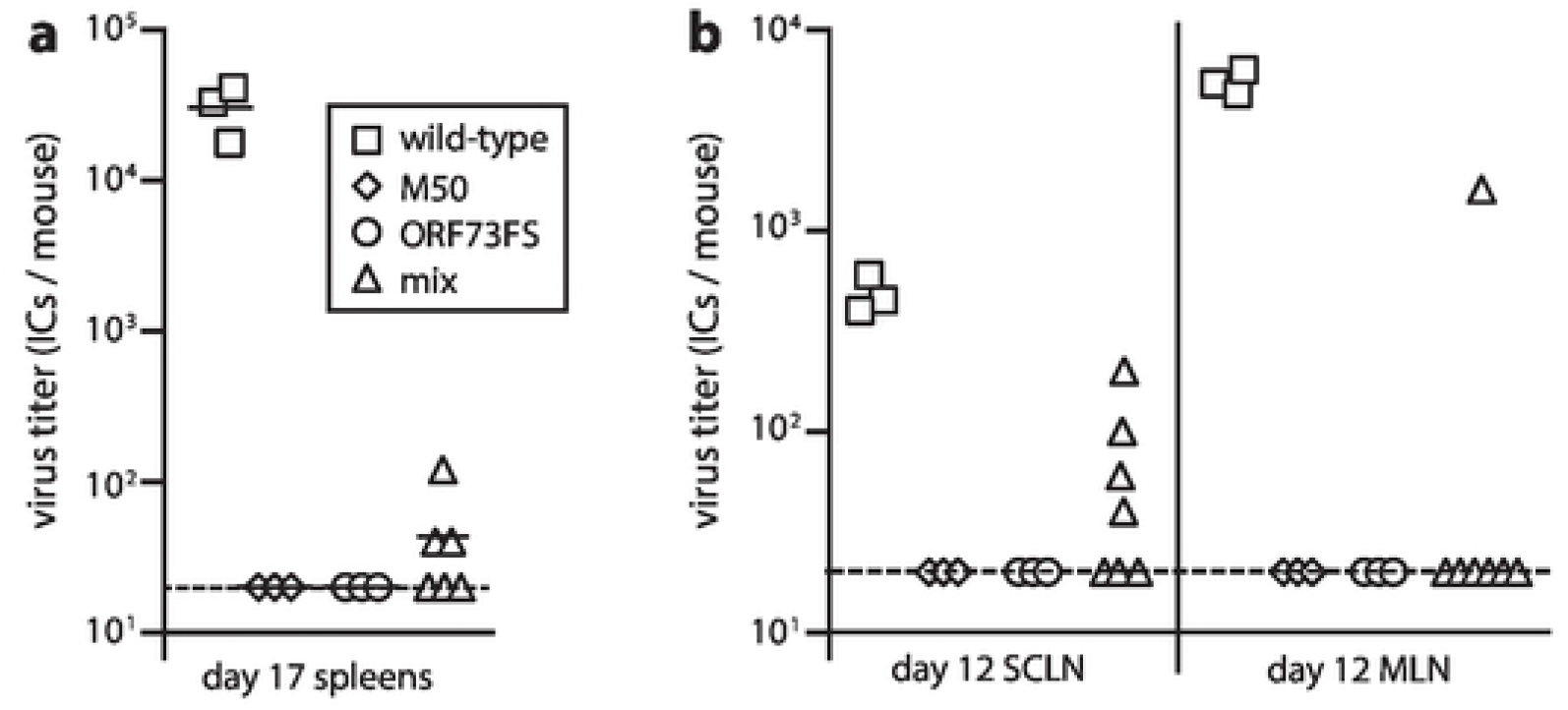
Little evidence of MuHV-4 recombination after lung co-infection. **a.** C57BL/6 mice were infected in the lungs (10^4^ p.f.u. in 30μl under anaesthesia) with wild-type, M50 or ORF73FS viruses, or a 1:1 M50 / ORF73FS mix. Spleens were infectious centre assayed for latent virus 17 days later. Symbols show individual mice, bars show means. The dashed line is the detection limit. **b.** Mice were infected as in **a.** 12 days later, SCLN and MLN were infectious centre-assayed for latent virus.

Comparison with nasal infection (Fig.2b) indicated less efficient rescue via the lungs, both in number of mice with detectable latent infection and in their titers. However direct comparison is difficult between mice infected by different routes. Also, while nasal infection does not reach the lungs, lung inocula must navigate the nose, so the low level splenic latency seen in Fig.4a might have come from contaminating olfactory infection. To address these points we assayed separately the superficial cervical (SCLN) and mediastinal lymph nodes (MLN) of co-infected mice, for latent virus after 12 days (Fig.4b). While lung and olfactory infections both spread via lymph nodes, the SCLN drain the nose whereas the MLN drain the lungs. Mononucleosis-associated systemic viral spread reaches far lymph nodes, but early infections reflect primary replication in peripheral sites. Specifically, low dose day 12 olfactory infections spare the MLN, so infection routes can be distinguished. At this time 4/7 mice had a low level of SCLN latency, and 1/7 mice had MLN latency. Wild-type controls showed significantly more MLN than SCLN infection, matching the more extensive lytic infection of lungs; and the sole positive MLN of the co-infected cohort had a correspondingly higher titer than any SCLN. Thus, we could be confident of not missing recombinational rescue in the lungs. It seemed likely that the low-level day 17 spleen infections in Fig.4a resulted from virus caught inadvertently in the upper respiratory tract, while lung co-inocula rarely recombined: 1/7 mice, versus 11/12 for deliberate olfactory infection in Fig.2b (p<0.002).

### Recombination via olfactory infection but not lungs applies also to MCMV

To explore whether recombinational rescue was specific to MuHV-4, we tested MCMV, co-infecting BALB/c mice with mutants lacking M33 or M78 (Fig.5). MCMV needs both these G protein-coupled receptor homologues to successfully colonize the salivary glands [39, 40]. Salivary gland plaque assays at 18 days post-lung inoculation were positive only for the wild-type control (Fig.5a). However at day 18 after nasal co-infection, salivary glands were positive for all co-infected mice, and for no M33^-^ or M78^-^ single controls (Fig.5b).

**Figure 5.**
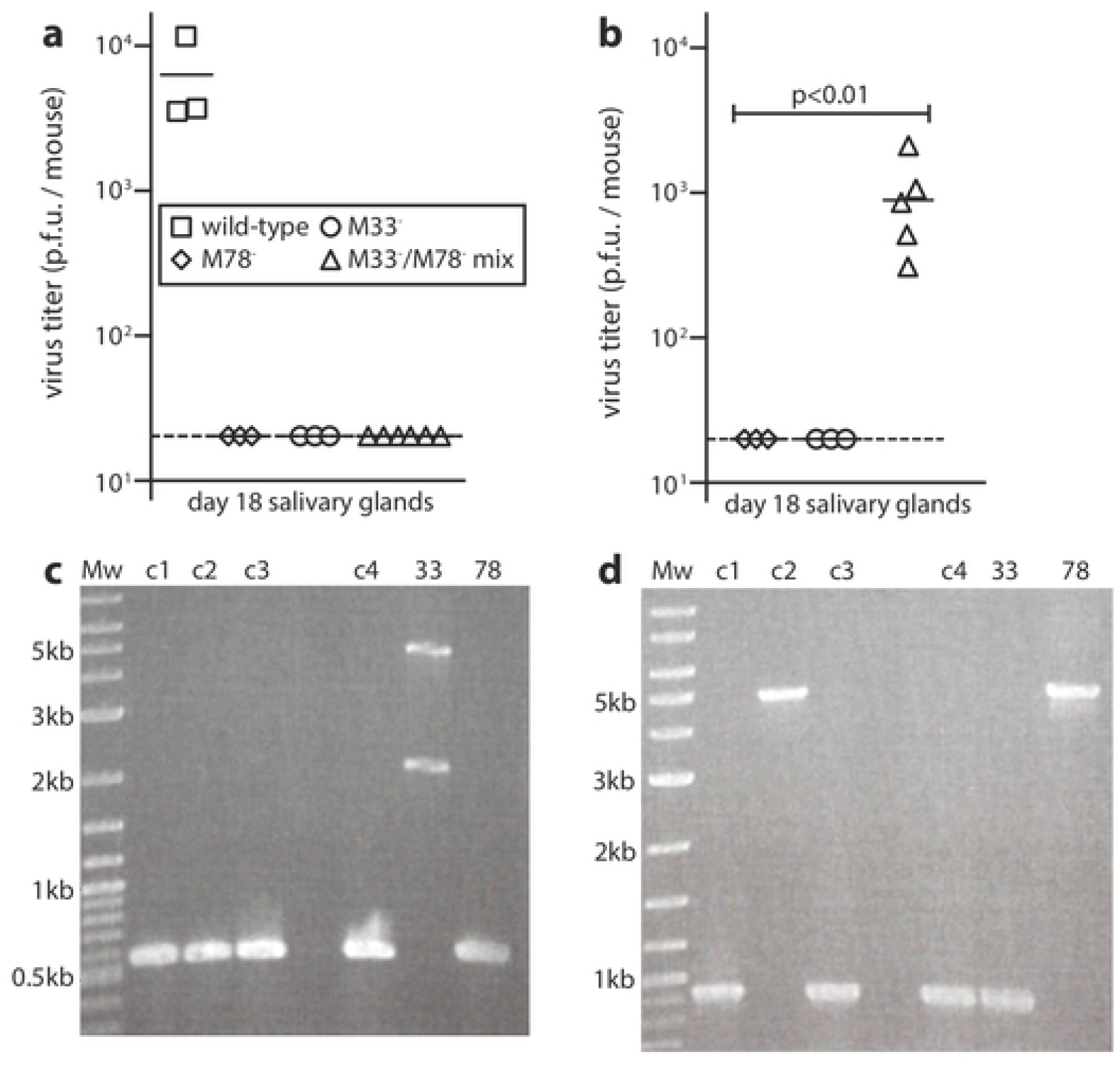
Rescue of MCMV mutants after co-infection in the nose but not the lungs. **a.** BALB/c mice were infected in the lungs (10^4^ p.f.u.) with wild-type, M33^-^ or M78^-^ MCMV, or a 1:1 M33^-^ / M78^-^ mix. 18 days later, salivary glands were plaque-assayed for infectious virus. Symbols show individual mice, bars show means. The dashed line is the detection limit. Only wild-type infection yielded recoverable virus. **b.** BALB/c mice were infected nasally (10^5^ p.f.u.) with M33^-^ or M78^-^ MCMV, or a 1:1 M33^-^ / M78^-^ mix. 18 days later, salivary glands were plaque-assayed for infectious virus. Symbols show individual mice, bars show means. Only the mixed infection yielded recoverable virus. **c.** DNA of virused cloned from mixed infection salivary glands in **b** was checked for M33 mutation by PCR. 33 and 78 are the M33^-^ and M78^-^ input viruses. The predicted wild-type band is 572bp. The upper 33 sample band corresponds to the expected size for the lacZ cassette insertion of the mutant (4.4kb). The source of the lower (2.2kb) 33 sample band is unclear, but the 33 sample is clearly free of wild-type DNA, and no recovered clone shows a mutant M33 locus. **d.** The same DNA samples were checked for M78 mutation. Clone 2 appeared to be a parental M78^-^ virus. The other 3 clones had a wild-type M78 locus, indicating recombination.

Viruses cloned from the salivary glands of 4/4 nasally co-infected mice showed a wild-type M33 locus by PCR (Fig.5c). One clone retained a mutant M78 locus (Fig.5d), perhaps because M78-deficient viruses do not always completely lack salivary gland infection [41, 42]. Nonetheless recombination was evidently the main route of rescue. Overall, MCMV like MuHV-4 showed recombinatorial rescue after olfactory infection but not after lung infection.

### Recombination occurs early in olfactory infection

MuHV-4 lung and olfactory infections show similar systemic spread, both being brought by dendritic cells to B cells, while lung and olfactory MCMV infections both spread via dendritic cell recirculation [40, 43]. Thus inefficient recombination via the lungs argued that it occurred early in olfactory infection. Supporting this supposition, infectious centre assays detected more latent MuHV-4 in SCLN at 8 days after nasal M50 / ORF73FS co-infection than after single infections (Fig.6a). The wide spread of co-infection titers at this early timepoint made its yield not significantly more than M50 alone. Therefore we looked further for evidence of recombination by inoculating virus clones from SCLN into the lungs of naive mice (Fig.6b). At day 14, clones recovered from M50 or ORF73FS single inoculations yielded no infectious centres, while all bar one of those from mixed inoculations - presumably a parental M50 virus - had titers indistinguishable from wild-type. Therefore olfactory MuHV-4 recombined before leaving lymph nodes.

**Figure 6.**
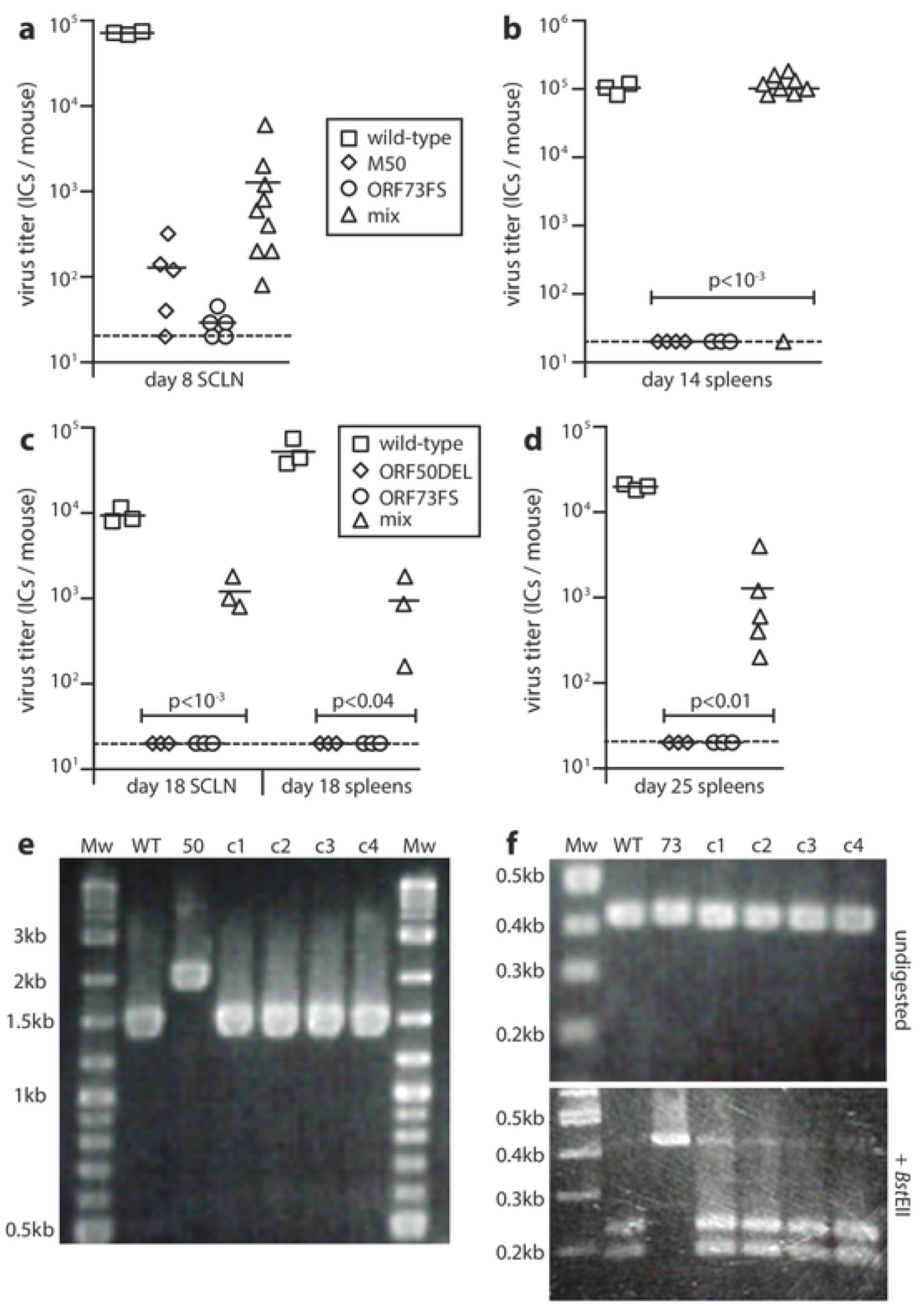
Early recombinational rescue of MuHV-4 after nasal co-infection. **a.** C57BL/6 mice were infected nasally (10^5^ p.f.u.) with wild-type, M50 or ORF73FS MuHV-4, or a 1:1 M50 / ORF73FS mix. 8 days later, latent virus in SCLN was infectious centre-assayed. Symbols show individual mice, bars show means. The dashed line is the detection limit. Mixed infections gave higher titers than either component single infection, although the wide spread meant that as a group they were not significantly higher than M50 alone. **b.** Viruses cloned from SCLN for each mouse in **a** were given i.n. to naive mice under anaesthesia to inoculate the lungs (10^3^ p.f.u. in 30μl). One clone from each positive mouse was given to one naive mouse. 14 days later spleens were infectious centre-assayed for latent infection. Symbols show individual mice, bars show means. **c.** C57BL/6 mice were infected nasally (10^5^ p.f.u. in 5μl) with wild-type, ORF50DEL or ORF73FS MuHV-4, or a 1:1 ORF50DEL / ORF73FS mix. 18 days later, SCLN and spleens was infectious centre-assayed for latent virus. Symbols show individual mice, bars show means. Mixed infections gave significantly higher titers than either component single infection. **d.** Mice infected as in **c** were infectious centre-assayed for splenic virus after 25 days. Mixed infections gave significantly higher titers than either component single infection. **e.** DNA of viruses cloned (c1-c4) from mixed infection spleens in **d** was PCR amplified across the ORF50 deletion site (primers p3 and p4 in Fig.1a). The PCR products were resolved by agarose gel electrophoresis and stained with ethidium-bromide. WT and 50 show wild-type and ORF50DEL input viruses. Mw = molecular weight markers. DNA sequencing of cloned virus PCR products confirmed identity with the wild-type. **f.** DNA from the mixed infection clones in **e** was PCR amplified across the ORF73 frameshift site (primers p5 and p6 in Fig.1a). The PCR products were digested or not with *Bst*EII, resolved by agarose gel electrophoresis, and stained with ethidium-bromide. WT and 73 are the wild-type and ORF73FS input viruses. As in Fig.2d, as c1-c4 were cloned prior to analysis, the minor residual 0.43kb band after incubation with *Bst*EII digestion presumably reflected incomplete digestion rather than mutant DNA. Sequencing of the undigested PCR products confirmed identity with the wild-type.

We recovered no recombinant viruses from co-infected noses. This likely reflected that without the selective pressure of having to establish latency, the proportion of recombinants remained low; and as M50 MuHV-4 out-replicates the wild-type *in vitro* [29], it predominated when producing stocks from a mixed population. Therefore to position more precisely the place of recombination, we co-infected mice with ORF73FS and ORF50DEL MuHV-4 (Fig.6c). The latter mutant lacks all lytic replication, due to a large deletion in ORF50 (Fig.1a). *In vitro* it must be propagated in complementing ORF50^+^ cells [20]; *in vivo* it cannot spread further than the first encountered cells [25]. Nonetheless, spleens and SCLN at 18 days after nasal co-infection yielded recoverable virus for 3/3 mice, and spleens at after 25 days did so for 5/5 mice. Therefore co-infection must have occurred in the initially infected cells. ORF73FS MuHV-4 could in principle complement ORF50DEL *in trans*, but an ongoing need for co-infection would render this less efficient than recombination, and viruses cloned from day 25 spleens of 4/4 co-infected mice showed wild-type ORF50 and ORF73 loci by PCR (Fig.6d, 6e). DNA sequencing confirmed identity with the wild-type.

### Recombination does not require simultaneous co-infection

ORF50DEL MuHV-4 cannot replicate lytically but can establish a latent infection of olfactory neurons and sustentacular cells [22]. We reasoned that it might remain accessible to recombination with a super-infecting virus, and tested this by infecting mice nasally with ORF50DEL MuHV-4, then 5 days later giving the same mice nasal ORF73FS MuHV-4 (Fig.7). We assayed spleens for recoverable virus another 18 days later. 5/6 co-infected mice showed splenic infection, while no singly infected controls did so. Therefore latent infection supplied a sufficient substrate for subsequent olfactory rescue.

**Figure 7.**
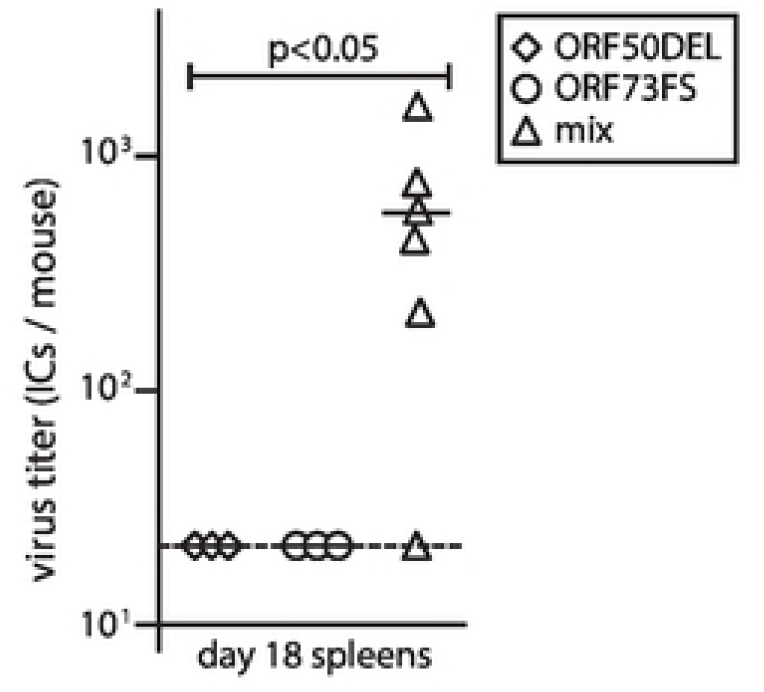
Recombinational rescue after serial mixed infection. **a.** C57BL/6 mice were infected nasally (10^5^ p.f.u.) with ORF50DEL MuHV-4. 5 days later they were infected nasally (10^5^ p.f.u.) with ORF73FS MuHV-4. Controls were given either just ORF50DEL or just ORF73FS. 18 days after ORF73FS infection, spleens were infectious centre-assayed for latent virus. Symbols show individual mice, bars show means. The dashed line shows the detection limit. Serial mixed infection yielded significantly more recoverable virus than either single infection.

### A possible site of olfactory recombination

Olfactory cells far out-number inoculated virions, and initially few are infected [22]. Dilution should reduce recombination. However the distribution of early olfactory infection is not uniform. A conspicuously common site is the respiratory / olfactory epithelial border (Fig.8). Anatomical complexity [44] makes this border hard to check completely, but retrospective review identified its involvement in 23/30 early infections (1-3 days post-inoculation). Generally noses were examined only until at 50-100 infected cells were found, so involvement of the border was likely under-estimated. Frequent infection initiation here would explain the recombinatorial rescue of replication-deficient virus.

**Figure 8.**
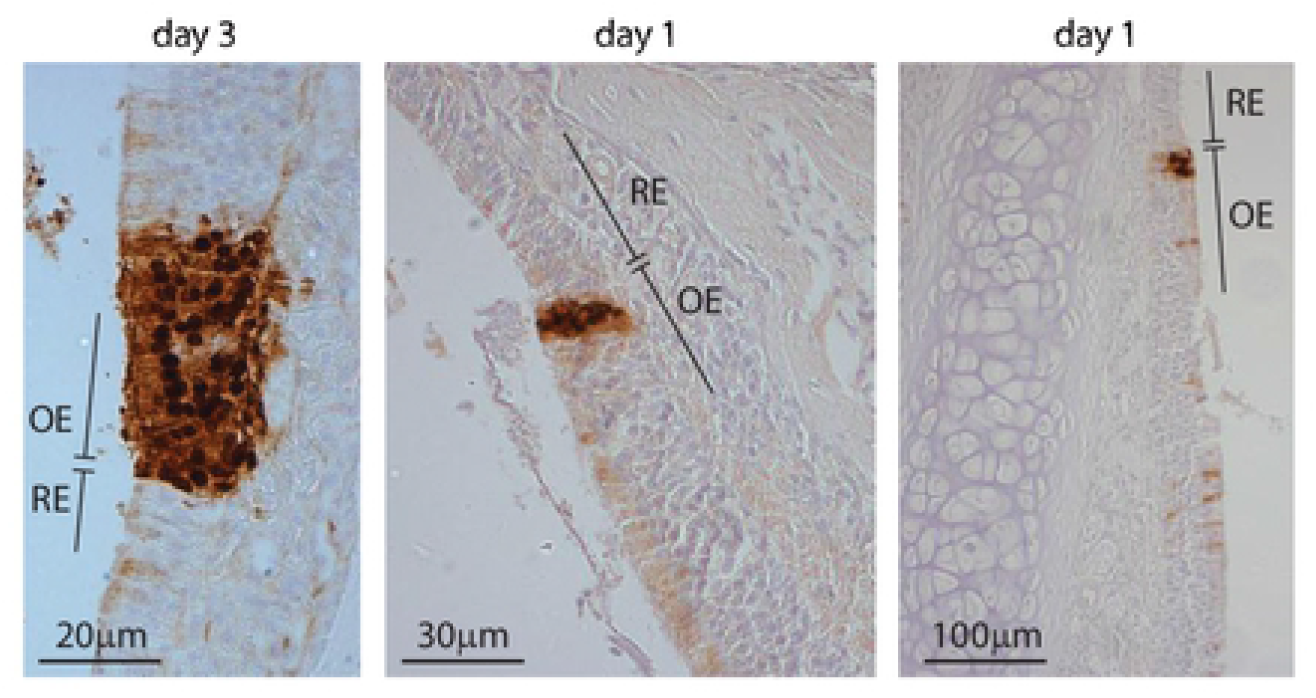
Infection at the olfactory / respiratory border. C57BL/6 mice were infected nasally with wild-type MuHV-4 (10^5^ p.f.u.). 1-3 days later, nose sections were stained for viral lytic antigens with a polyclonal rabbit serum (brown), and counter-stained with haemalum (blue). OE = olfactory epithelium, RE = respiratory epithelium. Representative sections show foci of infection on the olfactory side of the olfactory / respiratory border.

## Discussion

Herpesviruses interact with polymorphic host genes, and so encounter environmental change in each infection. Unlinking viral genes can generate new combinations to cope. Host genes mix in meiosis; viruses must count on co-infecting cells. While herpesviruses show historic recombination, whether this is occasional and mostly old, or whether viral genes reassort routinely has been obscure. We showed that for both MCMV and MuHV-4, recombination after olfactory entry routinely rescues replication defects. It provided repair where point mutations did not, and was efficient enough to overcome immune priming by the parent viruses.

Rescue of replication-deficient MuHV-4 implied co-infection of the first entered cells. The olfactory cells sit above and behind the respiratory surface, near the cribriform plate, which their axons broach to reach the brain [44]. Thus inhaled droplets reach the respiratory epithelium first, and surface tension likely captures them onto mucus. Consistent with this concept, inhaled virions appear immediately on the respiratory epithelium then disappear from here, while on the olfactory epithelium they accumulate and infect [22]. Respiratory epithelial cells are not infected because they lack apical heparan. Instead their cilial beating bears mucus-bound virions backwards. Hence infection commonly occurred close to the respiratory / olfactory border. Such concentration of captured virions should promote co-infection. It is probably more pronounced still in natural transmission, as negligible volume would then minimize nasal fluid flow. Cilial clearance of captured virions would explain further why olfactory infections fail to involve the lungs [20, 26, 27], as bronchial cilial push their mucus upwards to be swallowed.

MuHV-4 binds to olfactory neuronal cilia. Yet in adult mice it infects mostly adjacent sustentacular cells, which accumulate virions on their apical microvilli [22]. The neuronal cilia are too fine to transport virions internally, so retrograde cilial transport may move bound virions externally across the olfactory mucus. Sustentacular microvilli could then collect them. Mutant rescue via the lungs was rare, despite more lytic replication happening here. MuHV-4 lung entry involves macrophages capturing virions from alveolar epithelial cells [45], reminiscent of sustentacular cells capturing them from olfactory cilia. However inoculated virions are diluted among many more alveoli [45, 46]. Thus there may be little co-capture, and whether infection can spread between alveoli is far from clear.

Limited rescue via the lungs argued against frequent recombination down-stream of the olfactory epithelium, as subsequent spread seems similar between these routes - via dendritic cells to B cells. Once a tissue barrier is breached immunity begins, so infection must spread fast, implying selection for divergence over convergence. Down-stream dilution would explain how recombinants escaped the dominant negative impact of M50 on latency. While the latency defects of M50 and ORF73FS MuHV-4 made recombination in spleens impossible to assay, spread here recapitulates the myeloid / lymphoid virus relay into SCLN [24], via reactivation from B cells [47] then virion capture and transfer by marginal zone macrophages [25]. Thus it provides no obvious point of convergence. Ultimately, many reactivations pool progeny virions in saliva, but recombination requires that these virions co-infect. The obvious site for this is in new hosts.

EBV carriers can co-shed virus strains acutely [48], but long term infections seem to select single dominant strains [49]. So if co-infection depended completely on carriers co-shedding - if it takes a co-infection to make a co-infection - how could could it remain common? There must also be a means to mix serially infecting viruses. MuHV-4 vaccine assays argue that anti-viral immunity inhibits super-infection [46], even if the immunizing virus persists poorly [35, 36]. However establishing effective immunity takes time. The antibody response to MuHV-4 does not reach its peak for at least 3 months [50]; EBNA-specific antibodies appear may months after EBV acquisition, indicating immunity is still maturing; and EBV loads remain elevated for at least as long [51]. Thus an infant infected by a single maternal strain could acquire more from other carers [52] while still susceptible to super-infection.

Primary olfactory infection is soon suppressed [20, 26, 27]. Yet the proximity of mouth and nose means infected hosts inevitably inhale some of their own shed salivary virus. So carriers may continue to display their resident viruses for recombination. Latency in olfactory epithelial cells provides another means to meet. Olfactory entry by herpesviruses depends on heparan binding [22]. As such binding is widely shared, other viruses might also meet at the respiratory / olfactory border. The inherent dilution of transmission presents a problem for adeno-associated virus, to find an assisting adenovirus in each new host; but as both bind heparan [53, 54], adeno-associated virus could use latency in the olfactory epithelium to ambush its essential aide.

## Materials and Methods

### Mice

C57BL/6 or BALB/c mice were infected intransally (i.n.) when 6–14 weeks old, either in 30μl under isoflurane anaesthesia to reach the lungs (10^4^ p.f.u.), or in 5μl without anaesthesia (10^5^ p.f.u.), to infect only the upper respiratory tract. A higher dose was used for upper respiratory tract infection, as most of the inoculum is swallowed [39]. Statistical comparison was by Student’s 2-tailed unpaired *t*-test unless stated otherwise.

### Ethics statement

Experiments were approved by the University of Queensland Animal Ethics Committee in accordance with the Australian code for the care and use of animals for scientific purposes, from the Australian National Health and Medical Research Council (projects 391/15, 479/15, 196/18, 207/18).

### Cells and Viruses

BHK-21 fibroblasts (American Type Culture Collection (ATCC) CCL-10), NIH-3T3 cells (ATCC CRL-1658) and NIH-3T3-ORF50 cells [20] were grown in Dulbecco’s modified Eagle’s medium (Gibco) with 2mM glutamine, 100IU/ml penicillin, 100μg/ml streptomycin and 10% fetal calf serum (complete medium). Murine embryonic fibroblasts were grown in similarly supplemented minimal essential medium. MuHV-4 with a frameshift inactivating ORF73 [32], with an additional deletion of 10kb from the genome left end, that includes ORFs M1, M2, M3, M4 and 4 [37], with a deletion of ORF50 exon 2 [20], or with an MCMV IE1 promoter fragment inserted in the 5’ untranslated region of ORF50 [30] are described. ORF50DEL MuHV-4 was grown and titered in NIH-3T3-ORF50 cells. Other MuHV-4 derivatives were grown and titered in BHK-21 cells. MCMV mutants with a β-galactosidase expression cassette disrupting M33 [40] or M78 [41] were derived from K181 strain Perth, which was used as the wild-type. All were grown in NIH-3T3 cells. Infected cell supernatants were cleared of debris by low speed centrifugation (200 x *g*, 5 min). Cell-free virions were then concentrated by ultracentrifugation (35,000 x *g*, 1.5h) and stored at −80°C.

### Infectivity assays

Infectious virus was quantified by plaque assay. For MuHV-4 [55], virus stocks or freeze-thawed tissue homogenates dilutions were incubated with BHK-21 cells (2h, 37°C), overlaid with complete medium / 0.3% carboxymethylcellulose, cultured for 4 days, fixed with 1% formaldehyde and stained with 0.1% toluidine blue for plaque counting. MCMV was titered similarly but on murine embryonic fibroblast monolayers [41], and was adhered by centrifugation (500 x *g*, 30 min) before discarding the inoculum. Total recoverable MuHV-4 (latent plus pre-formed infectious virus) was quantified by infectious centre (i.c.) assay [55]. Freshly isolated lymph node or spleen cells were layered onto BHK-21 cell monolayers then cultured and processed as for plaque assays. For parallel assays of pre-formed infectious virus, samples were first frozen and thawed.

### Viral Genome Quantitation

MuHV-4 genomic coordinates 4163-4308 (M2 ORF; primers 4163-4183 and 4288-4308) were amplified by PCR (Rotor Gene 3000, Corbett Research) from 50ng DNA (Nucleospin Tissue kit, Macherey-Nagel). PCR products quantified with Sybr green (Thermo Fisher Scientific) were compared to a standard curve of cloned template amplified in parallel, and distinguished from paired primers by melting-curve analysis. Correct sizing was confirmed by electrophoresis and ethidium bromide staining. Cellular DNA in the same samples was quantified by parallel amplification of a β-actin gene fragment.

### Virus genotyping

M50 MuHV-4 has the proximal 416bp of the murine cytomegalovirus IE1/IE3 promoter inserted at genomic coordinate 66718, between the ORF50 transcription (66642) and translation start sites (66760). To detect this insert we amplified across genomic coordinates 66580-66848, using primers 5’-cacattatcccacaatgtgctgc and 5’-gaaatactgatctgtctgcgtgg. This gave a 268bp wild-type product and a 684bp M50 product. ORF50DEL MuHV-4 has genomic coordinates 67792-69177 deleted from ORF50 exon 2 (67661-69376). In place is ligated a 1961bp fragment comprising the firefly luciferase coding sequence in frame with ORF50 (1672bp) and an SV40 polyadenylation signal (289bp). We amplified across genomic coordinates 67672-69240, using primers 5’-gatcgaagcaggtctacttgag and 5’-tcagcagtgtcctggtttgcc, to give a 1569bp wild-type band or a 2145bp mutant band. ORF73FS MuHV-4 has a 5bp insert at genomic coordinate 104379 in ORF73 (104869-103925). This disrupts a *Bst*EII restriction site. We amplified across genomic coordinates 104152-104580, using primers 5’-ttcacagtaggccaagacaacc and 5’-ccaccatcaccagatgttgatg. This gave a 428bp wild-type product, cut by *Bst*EII into 228bp and 200bp fragments; or a 433bp mutant product not cut by *Bst*EII. DNA sequencing of PCR products was performed at the Australian Genome Research Facility (St.Lucia, Queensland). To identify MCMV M33 disruption we amplified viral DNA with primers 5’-gtggtgctgacgacgcagctgctg and 5’-gtgtggctgcgcctgcggtacgag for a wild type band of 572bp. The M33^-^ mutant has a 3.8kb HCMV IE1 promoter - lacZ - poly A cassette inserted without deletion to give a 4.4kb band. To identify M78 disruption we amplified viral DNA with primers 5’-tcgtctgcccctctaaggtca and 5’-cagacggtggggatcttgtcg for a wild type band of 919bp. The M78^-^ mutant has an equivalent 3.8kb cassette insertion to give a 4.7kb band.

### Immunostaining tissue sections

Organs were fixed in 1% formaldehyde / 10 mM sodium periodate / 75 mM L-lysine (18h, 4°C). Noses were then decalcified in 150mM NaCl / 50mM TrisCl pH 7.2 / 270mM EDTA for two weeks at 23°C, changing the solution every 3 days, then washed twice in PBS. Samples were dehydrated in graded ethanol solutions, embedded in paraffin, and cut with a microtome. Sections were then de-waxed in xylene, rehydrated, washed 3x in PBS and air-dried. Endogenous peroxidase activity was quenched in PBS / 3% H_2_O_2_ for 10min. Sections were blocked (1h, 23°C) with 0.3% Triton X-100 / 5% normal donkey serum, and incubated (18h, 4°C) with anti-MuHV-4 rabbit pAb, which recognizes a range of virion proteins by immunoblot, including the products of ORF4 (gp70), M7 (gp150), and ORF65 (p20) [55]. Sections were additionally blocked with Avidin/Biotin Blocking Kit (Vector Laboratories). Detection was with biotinylated goat anti-rabbit IgG pAb (1h, 23°C, Vector Laboratories), Vectastain Elite ABC Peroxidase system, and ImmPACT diaminobenzidine substrate (Vector Laboratories). Stained sections were counterstained with Mayer’s Hemalum (Merck), dehydrated and mounted in DPX (BDH).

